# Structural determinants of the direct inhibition of GIRK channels by Sigma-1 receptor antagonist

**DOI:** 10.1101/2023.11.07.566128

**Authors:** Chang Liu, I-Shan Chen, Michihiro Tateyama, Yoshihiro Kubo

## Abstract

G-protein-gated inward rectifier K^+^ (GIRK) channels play critical roles in the regulation of the excitability of cardiomyocytes and neurons and include GIRK1, GIRK2, GIRK3 and GIRK4 subfamily members. BD1047 dihydrobromide (BD1047) is one of the representative antagonists of multi-functional Sigma-1 receptor (S1R). In the analysis of the effect of BD1047 on the inhibition of Gi-coupled receptors by S1R using GIRK channel as an effector, we observed that BD1047 directly inhibits GIRK current even in the absence of S1R. Thus, we aimed to clarify the effect of BD1047 on GIRK channels and its structural determinants. By electrophysiological recordings in *Xenopus oocytes*, we observed that BD1047 directly inhibited the current of GIRK channels, producing a much stronger inhibition of GIRK4 channels compared to GIRK2. It also inhibited the ACh-induced native GIRK current in isolated rat atrial myocytes. Chimeric and mutagenesis studies of GIRK2 and GIRK4 combining with molecular docking analysis, demonstrate the importance of the Leu77 on the proximal N-terminal cytoplasmic region of GIRK4 for the inhibition by BD1047. The activator of GIRK channel, ivermectin, competed with BD1047 at Leu77 on GIRK4. This study provides us with a novel inhibitor of GIRK channels and information for developing pharmacological treatments for GIRK4 associated diseases.

**Key points:**

- Sigma-1 receptor antagonist, BD1047, directly inhibits the current of GIRK channels. It strongly inhibits GIRK4 channels but only weakly inhibits GIRK2.
- BD1047 inhibits the ACh-induced GIRK current in isolated rat atrial myocyte.
- Leu77 on the proximal N-terminal of GIRK4 is essential for the inhibition by BD1047.
- Binding of BD1047 adjacent to Leu77, but not to Leu77Ile, was confirmed by molecular docking analysis.
- The activator of GIRK channel, ivermectin, competes with BD1047 at Leu77 on GIRK4.

## Introduction

The G-protein-gated inward rectifier K^+^ (GIRK) channel is known to be activated by the G_βγ_ subunit released from stimulated G_i_-coupled G protein-coupled receptors (GPCRs) (Logothetis *et al*., 1987; Clapham & Neer, 1993) and it is involved in wide variety of physiological functions, such as the regulation of excitability of neurons and the heat rate (Hibino *et al*., 2010). Given these important roles, inhibitors and activators of GIRKs have been broadly investigated. In addition to G_βγ_ subunit, GIRK channel were shown to be directly activated by Na^+^, PIP_2_, N-alcohols, Naringin, ML297, ivermectin (IVM) and 18-β glycyrrhetinic acid (Huang *et al*., 1998; Ho & Murrell-Lagnado, 1999; Yow *et al*., 2011; Bodhinathan & Slesinger, 2013; Wydeven *et al*., 2014; Chen & Kubo, 2018; Chen *et al*., 2023). Direct inhibitors were also identified including non-selective pore blockers such as Ba^+^ and Cs^+^, selective inhibitor tertiapin-Q, antipsychotic drugs such as thioridazine, clozapine, antidepressant drugs like imipramine and fluoxetine, and antihistamine like terfenadine (Kobayashi *et al*., 2000; Kanjhan *et al*., 2005; Kobayashi *et al*., 2011; Chen *et al*., 2019; Jeremic *et al*., 2021).

There are four members of the GIRK family (GIRK1-4) which can form functional homo- and hetero-tetramers and are expressed in a variety of native tissues. For example, heteromeric GIRK1/2 channels were identified in brain, whereas, GIRK1/4 and homomeric GIRK4 channels are mainly expressed in atrial myocytes (Kubo *et al*., 1993; Lesage *et al*., 1994; Krapivinsky *et al*., 1995; Corey & Clapham, 1998). Previous studies have suggested that GIRK4 channel might be a promising therapeutic target for atrial fibrillation (AF). Because the induction of AF is inhibited in GIRK4 knock out mice (Kovoor *et al*., 2001), and the acetylcholine (ACh)-regulated potassium current, GIRK1/4, constitutively activate in atrial myocytes from AF patients (Dobrev *et al*., 2005). Thus, the identification of GIRK4 inhibitors may provide information relevant for drug design targeting GIRK4 related cardiac arrythmia.

Sigma-1 receptor (S1R) is a multimodal chaperone protein located chiefly at the mitochondrion-associated endoplasmic reticulum (ER) membrane (MAM) at rest. It can translocate to various regions of the cell such as plasma membrane (PM) or ER-PM junction under cellular stress or the application of its ligands (Su *et al*., 2016) and has been implicated in a diverse array of pathophysiological conditions including drug addiction, Parkinson’s disease, Alzheimer’s disease and amyotrophic lateral sclerosis (Kourrich *et al*., 2012; Su *et al*., 2016). BD1047 dihydrobromide (BD1047) is one of the representative antagonists of sigma-1 receptor (S1R) (Su *et al*., 2010) and in recent animal experiments was shown to be effective for the treatment of neuropathic pain through affecting the expression of S1R in the ipsilateral spinal cord dorsal of rat after chronic constriction injury (Skuza & Rogoz, 2006; Roh *et al*., 2008). BD1047 also plays a role in attenuating ethanol-induced neurotoxicity through modulating the function of endoplasmic reticulum (ER)-bound inositol triphosphate (IP_3_) and S1R in rat hippocampal, suggesting its possible use in alcohol induced disorder (Reynolds *et al*., 2016). It was reported that S1R is colocalized with G_i/o_-coupled muscarinic receptor M2 on the soma of motoneurons (Mavlyutov *et al*., 2010). Thus, it is possible that S1R regulates the function and/or expression of M2 receptor.

In our analysis of the inhibitory effects of S1R on GPCR signalling, we used GIRK channel as the effector of the G_i/o_ protein-coupled receptors. We tried to inhibit the function of S1R using BD1047, and unexpectedly observed that BD1047 directly inhibits the current of GIRK channel currents in the absence of S1R expression. Since this is a novel inhibitor of GIRKs, our aim was to identify the structural determinants involved in the binding interaction. By electrophysiological recordings in X*enopus* oocytes and isolated atrial myocytes of rat, together with mutagenesis studies and molecular docking, we have identified BD1047 as a more potent inhibitor of GIRK4 compared to GIRK2 channels and also shown that it inhibits ACh-induced native GIRK current in rat atrial myocytes. Leu77 within the proximal N-terminus (N-ter) of GIRK4 is critical for the inhibition by BD1047. We also observed the activator of GIRK channel, IVM, competes with BD1047 at Leu77.

## Methods

### Ethical approval

All animal experiments in this study were approved by the Animal Care Committee of the National Institutes of Natural Sciences (an umbrella institution of National Institute for Physiological Sciences, Japan), and were performed in accordance with its guidelines.

### Preparation of *Xenopus laevis* oocytes

Oocytes were isolated from *Xenopus laevis* (purchased from Hamamatsu Seibutsu Kyouzai, Hamamatsu, Japan) under anesthesia of 0.15% tricaine (Sigma-Aldrich) by surgery. The surgical operation on frog was performed on ice and an incision was made in the frog’s abdomen to remove the oocytes. Isolated oocytes were incubated in 2 mg mL^-1^ collagenase type I (Sigma-Aldrich) for 6 hours to remove the follicular membrane and incubated in frog Ringer’s solution containing 88 mM NaCl, 1 mM KCl, 2.4 mM NaHCO_3_, 0.3 mM Ca(NO_3_)_2_, 0.41 mM CaCl_2_, 0.82 mM MgSO_4_ and 15 mM HEPES with 0.1% penicillin-streptomycin at 17 °C. After the injection of 50 nL of cRNA, oocytes were incubated in frog Ringer’s solution for a further 3-4 days prior to electrophysiological recordings.

### Isolation of rat atrial myocytes

8-week female adult Wistar rats (180-200g) were purchased from Japan SLC, Inc. (Hamamatsu, Japan) and housed in square cages with wood shavings, two rats per cage, on a light–dark cycle (12:12 hr) at 23 ± 1°C and fed standard rat chow in the Animal Facility of the National Institute for Physiological Sciences (NIPS, Japan). Isolation of atrial myocytes were performed as preciously described (Chen *et al*., 2014; Chen *et al*., 2019). Rats were anaesthetized after the intraperitoneal (I.P.) injection of 10 mg·kg^−1^ xylazine hydrochloride and 100 mg·kg^−1^ thiopental sodium for 6-8 min following i.p. injection of 1,000 U·kg^−1^ heparin. Then successful anaesthesia was judged by the full disappearance of the pedal withdrawal reflex. The rats were sacrificed by taking out the heart from the thoracic cavity under anaesthesia and the heart was connected to a modified Langendorff perfusion system via the aorta. A cannula was inserted into the aorta with the continuously retrograde perfusion of the Ca^2+^-free Tyrode’s solution (137 mM NaCl, 5.4 mM KCl, 1 mM MgCl_2_, 10 mM glucose, and 10 mM HEPES, pH 7.4 with NaOH) containing 0.6 mg·ml^−1^ collagenase type II (Worthington) and 0.2 mg·ml^−1^ protease type XIV (Sigma-Aldrich) at 37°C for 30-35 min through coronary arteries via aorta. Then the KB solution (10 mM taurine, 10 mM oxalic acid, 70 mM K glutamate, 25 mM KCl, 10 mM KH_2_PO_4_, 11 mM glucose, 0.5 mM EGTA, and 10 mM HEPES, pH 7.3 with KOH) was used to wash the digested tissue. The atria was cut into pieces with scissors and transferred to a culture dish. Atrial myocytes were dissociated by gently shaking the cut tissue pieces using tweezer. Dissociated atrial myocytes were incubated in the KB solution and myocytes were re-seeded onto Poly-L-Lysine (PLL) coated glasses and used for electrophysiological recordings on the same day.

### Mutagenesis and cDNA and cRNA preparations

For experiments in *Xenopus* oocytes, cDNAs of mouse GIRK2 and rat GIRK4 were subcloned into pGEMHE. Mutations in GIRK2 and GIRK4 were introduced using the PfuUltra II Fusion HS DNA Polymerase kit (Agilent Technologies) and verified by DNA sequencing. After linearization of cDNA by restriction enzymes, complementary RNAs were transcribed using mMessage mMachine kit (Ambion). The amount of each cRNA injected per oocyte was as follows: for rat GIRK1 (8.3 ng), mouse GIRK2 (4.2 ng), rat GIRK4 (8.3 ng), bovine G-protein β1 (8.3 ng) and bovine G-protein γ2 (8.3 ng).

### Electrophysiological recording

Two-electrodes voltage clamp recordings were made from *Xenopus* oocytes 3-4 days post-injection and data were acquired using an oocyte clamp amplifier (OC-725C, Warner Instruments), a digital analogue converter (Digidata 1550A, Molecular Devices) and pCLAMP 10.5 software (Molecular Devices). Recording microelectrodes had a resistance of 0.1-0.5 MΩ when filled with 3 M potassium acetate with 10 mM KCl. The 96K^+^ extracellular (EC) recording solution contained 96 mM KCl, 3 mM MgCl_2_ and 5 mM HEPES (pH 7.5 with KOH) and the ND96 solution contained 96 mM NaCl, 2 mM KCl, 1 mM MgCl_2_, 1.8 mM CaCl_2_ and 5 mM HEPES (pH 7.5 with NaOH). All experiments were performed at 25-28 °C with continuous perfusion using a peristaltic pump (AC-2110 II, ATTA) and BD1047 was applied to the whole bath.

In the whole cell patch clamp recording from isolated rat atrial myocytes, data were acquired using a patch clamp amplifier (AXOPATCH 200B), a digital analogue converter (Digidata 1440A, Molecular Devices) and pCLAMP 10.7 software (Molecular Devices). Myocytes were attached to PLL coated glass coverslips 1-3 hours before recording and placed in the recording chamber, and membrane current were recorded under whole cell patch clamp using a glass micropipette with the access resistance of 3-5 MΩ when filled with 130 mM KCl, 5 mM Na_2_ATP, 4 mM MgCl_2_, 0.1 mM CaCl_2_, 3 mM EGTA, 0.2 mM GTP, and 10 mM HEPES (pH 7.4 with KOH). A Ca^2+^-free Tyrode’s solution was used until the whole cell patch clamp configuration was achieved to avoid a twitching motion of the cardiac myocytes. The solution was then switched to a 20K^+^ EC solution containing 115 mM NaCl, 20 mM KCl, 0.53 mM MgCl_2_, 1.8 mM CaCl_2_, 5.5 mM glucose, and 5.5 mM HEPES (pH 7.4 with NaOH). All experiments were performed at 25-28 °C and ligands (1 µM ACh, 3 µM tertiapin-Q (TPN-Q), BD1047) dissolved in 20K^+^ solution were applied by gravity flow using a muti-valve controller system (VC-8 Valve controller, Warner instruments) combined with SF-77B perfusion fast step (Warner Instruments) for inlet and washed out via a suction pipette by pressure for outlet.

### Molecular docking

Homology model of GIRK4 was generated based on the structure of GIRK2 (6XIT, RCSB protein data bank) using SWISS-MODEL (https://swissmodel.expasy.org/). The dimer form of GIRK2 was directly built using PyMOL software and the dimer form GIRK4 was made based on the homology model of GIRK4 using PyMOL software. The tetrameric structure of mutants, GIRK2 I82L and GIRK4 L77I, were built using SWISS-MODEL and then modified to dimer form by PyMOL. The chemical structure of BD1047 was downloaded from the PubChem (https://pubchem.ncbi.nlm.nih.gov/compound/BD-1047-Dihydrobromide). The dimer form of GIRK2, GIRK4, GIRK2 I82L or GIRK4 L77I with the structure of BD1047 were uploaded to SwissDock (http://www.swissdock.ch/) to obtain the docking results respectively. The computational docking results was presented with color marks using Chimera 1.16 software (https://www.cgl.ucsf.edu/chimera/olddownload.html).

### Chemicals

Oxotremorine-M (OXO-M) (Sigma-Aldrich) was dissolved in distilled water to make a 50 mM stock solution and further diluted in the bath solution for the final concentration 50 µM. BD1047 (Tocris Bioscience) was dissolved in distilled water to make a 50 mM stock solution and further diluted in the bath solution for the final concentrations from 0.1µM up to 500 µM. IVM (Sigma-Aldrich) was dissolved in DMSO to make a 10 mM stock solution and further diluted in the bath solution to 100 µM. ACh (Sigma-Aldrich) was dissolved in distilled water to make a stock solution and then diluted in the bath solution 1 µM. Sources of other materials are described in the relevant methods.

### Data analyses

Two-electrode voltage clamp data were analyzed by Clampfit 10.7 (Molecular Devices) and Igor Pro 5.0 (WaveMetrics) and are shown as mean ± SD from n single oocytes. The current amplitude of GIRK channel was recorded at -100mV before and after the application of BD1047. The inhibition percentage of GIRK current by BD1047 was calculated by the decrease in the current amplitude in the presence of 100 μM BD1047 from the current before its application at -100 mV (I_GIRK basal_ -I_GIRK BD1047 (+)_ / I_GIRK basal_), and the basal current amplitude after subtraction of the current in ND96 solution was normalized as 100%. The dose-inhibition relationship was analyzed by sequential application of various concentrations of BD1047. Using Igor, the data were fitted to a Hill equation: y = I_min_ + (I_max_ − I_min_) / (1 + (x/IC_50_) ^ (Hillslope)). To analyse the effects of IVM on the rate of BD1047’s inhibition, the maximal inhibited current amplitude was normalized as 1, the normalized inhibited current in temporal variation was calculated from the beginning to the end of the application of BD1047 (I_normalized BD inhibited current_ = (I_BD_-I_BD begin_)/(I_BD final_ -I_BD begin_)). Then data were fitted to exponential formula: 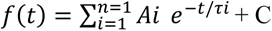 and tau of the inhibition was calculated by the fitting by Clampfit 10.7 (Molecular Devices).

In the patch-clamp experiments, all data were analyzed by Clampfit 10.7 (Molecular Devices) and Igor Pro 5.0 (WaveMetrics) and are shown as mean ± SD from n single atrial myocytes. To eliminate the influence of the desensitization of ACh-induced current, the maximal amplitude of ACh-induced GIRK current at -100 mV was normalized as 1 after the substraction of the basal current and the normalized current in the absence or presence of inhibitor (BD1047 or TPN-Q) was measured at the same time point (after application of TPN-Q or BD1047 for 27s) and then calculated by I_normalized current_ = (I_ACh-GIRK inhibitor (+) (27s)_ - I_basal_)/(I_ACh-GIRK(max)_ - I_basal_). The normalized current of 0 means there was no remaining ACh-induced GIRK current after the application of inhibitors.

All experiments were performed with the number of n ≥ 4 for each group. Statistical significances of differences were evaluated using Tukey’s multiple comparison tests following one-way ANOVA or unpaired *t*-test. The values of *P<*0.05 were judged to be statistically significant.

## Results

### BD1047 directly inhibits the current of GIRK channels in *Xenopus* oocytes

To examine the effects of BD1047 (**Fig. 1A**) on GIRK channels, *Xenopus* oocytes were used as an in vitro expression system and two-electrodes voltage clamp experiments were performed. We tested the effects of 100 µM BD1047 on different subtypes of GIRK channel, homomeric GIRK2 and GIRK4, and also on the heteromeric GIRK1/2 and GIRK1/4. Since homomeric GIRK4 channels show a small basal current (Krapivinsky *et al*., 1995; Kubo & Iizuka, 1996; Hibino *et al*., 2010), its physiological activator G_βγ_ subunits were always co-expressed together with GIRK4 in order to clearly record its current. The application of BD1047 decreased the current amplitude of GIRK1/2, GIRK2, GIRK1/4 and GIRK4 by 51.0 ± 1.8 %, 37.7 ± 9.8 %, 47.4 ± 4.9 % and 79.9 ± 4.6 % respectively, showing that BD1047 directly inhibited the current of GIRK channels, producing a much stronger inhibition of GIRK4 channels compared to GIRK2 (**Fig. 1B-F**). The concentration-response relationship for the effects of BD1047 on GIRK2 and GIRK4 were measured, GIRK2 was co-expressed with G_βγ_ subunits to ensure that the effect of BD1047 on GIRK channel is independent of G proteins **(Fig. 2A, 2B)**. The IC_50_ values were 75.4 ± 9.7 µM for GIRK2 and 17.4 ± 3.7 µM for GIRK4 (**Fig. 2C**). The Hill coefficient for GIRK4 was 1.4 ± 0.1 (**Fig. 2C**), suggesting that binding of one BD1047 molecule to GIRK4 is sufficient to inhibit the channel.

**Figure 1.**
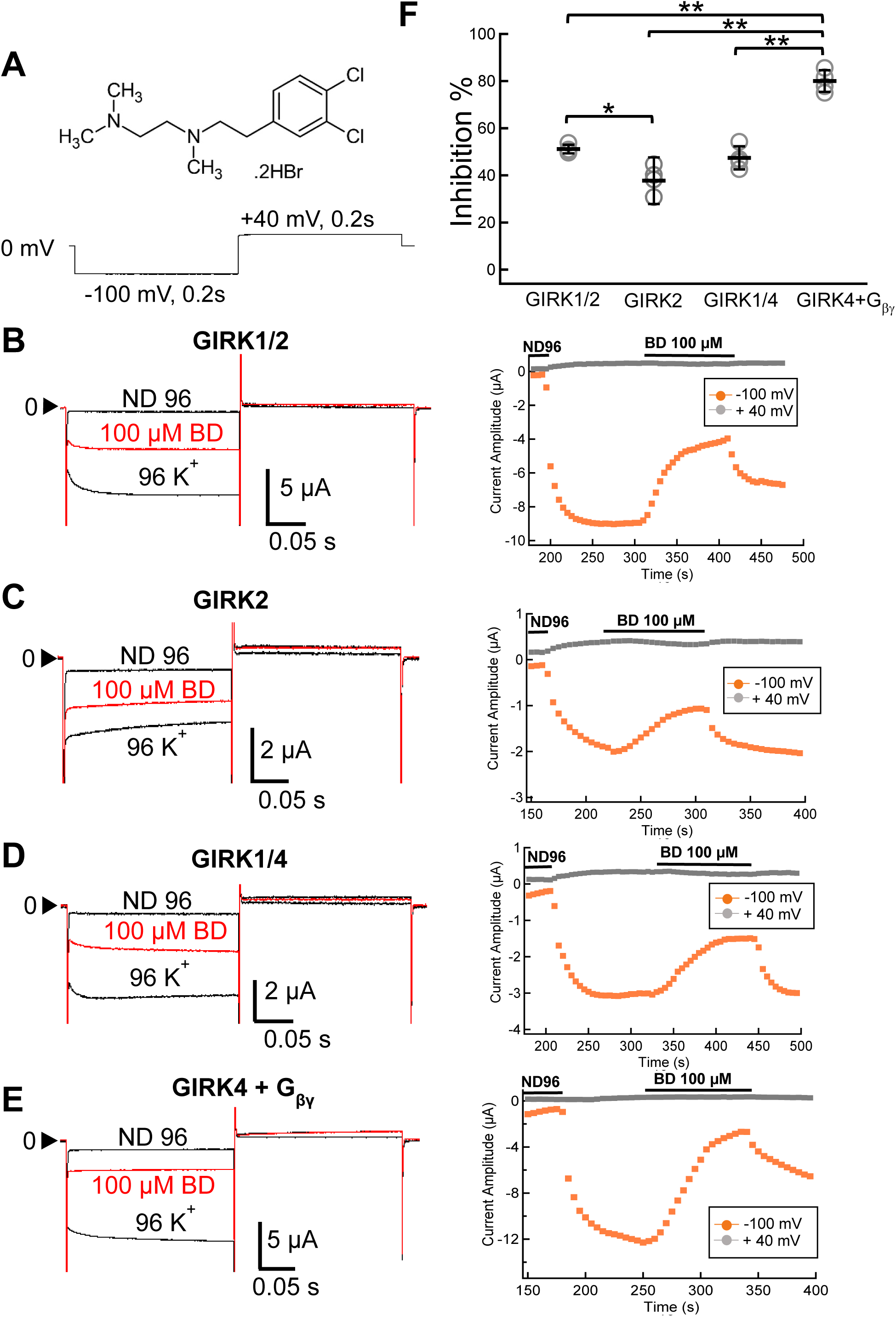
The effects of BD1047 on GIRK channels. **(A)** Chemical structure of BD1047 and the pulse protocol for recording. **(B-E)** Left: Representative current traces measured in *Xenopus* oocytes; Right: The timelapse changes of the current amplitudes at -100 mV (orange plots) and at +40 mV (gray dots) in ND96, 96K^+^ and 96K^+^ with 100 µM BD1047 solution in oocytes expressing (B) GIRK1/2, (C) GIRK2, (D) GIRK1/4 or (E) GIRK4+G_βγ_ channels. BD1047 was applied and washed out by constant bath perfusion using a peristatic pump. **(F)** Inhibition by 100 µM BD1047, of GIRK1/2, GIRK2, GIRK1/4 and GIRK4 channel currents. Data are mean ± SD (n = 5 for each); One way ANOVA followed by Tukey’s test, * indicates *P*<0.05; ** indicates *P*<0.01.

**Figure 2.**
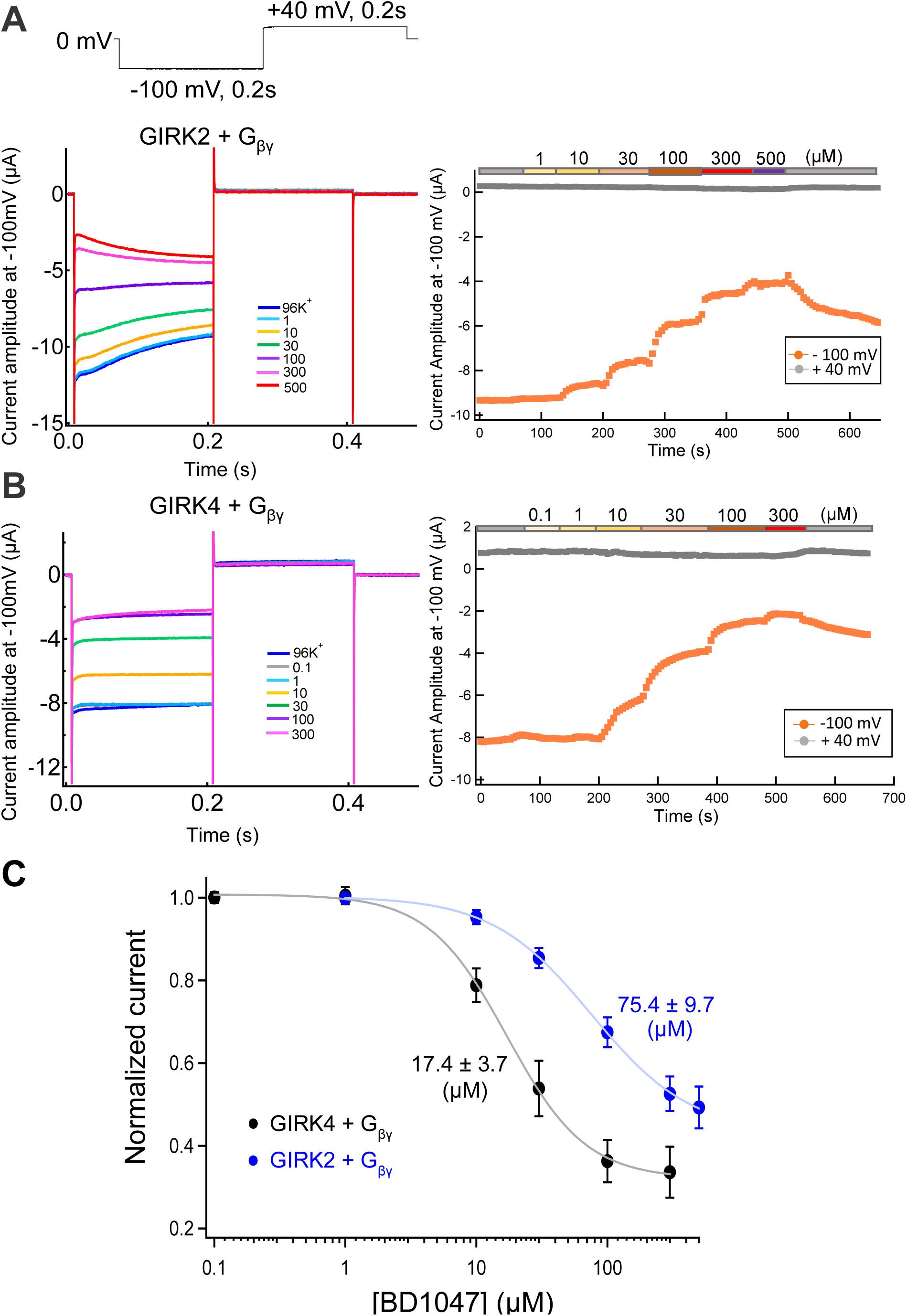
Dose inhibition relationships of BD1047 on GIRK2 and GIRK4 current. **(A-B)** Left: Representative current recordings in *Xenopus* oocytes evoked by the voltage protocol shown above. BD1047 ranging in concentration from 0.1 μM to 500 μM was applied to GIRK2 (A) and GIRK4 (B) channels. Right: the time courses of the current changes in various concentrations of BD1047 in oocytes expressing (A) GIRK2 or (B) GIRK4 channels. Orange dots indicate the recorded current amplitudes at -100 mV and gray dots indicate those at +40 mV. **(C)** Dose-inhibition relationships of BD1047 on GIRK2 (blue) and GIRK4 channel (black). Data are mean ± SD (n = 4-5) for each plot. IC_50_ is 75.4 ± 9.7 μM (GIRK2) and 17.4 ± 3.7 μM (GIRK4), respectively.

### BD1047 inhibits the ACh-induced GIRK current in rat atrial myocytes

GIRK1/4 channels are expressed in mammalian atrial myocytes (Hibino *et al*., 2010) and can be activated by ACh via the M2 muscarinic receptors to slow the heart rate (Krapivinsky *et al*., 1995; Hibino *et al*., 2010; Cui *et al*., 2021). We examined the effects of BD1047 on native GIRK currents in isolated rat atrial myocytes. The application of 1 µM ACh activated GIRK1/4 currents which slowly desensitized, a well-known phenomenon of ACh-induced GIRK current (Carmeliet & Mubagwa, 1986; Kurachi *et al*., 1987) (**Fig. 3B**). To eliminate the influence of the desensitization of ACh-induced current, the maximal amplitude of ACh-induced GIRK current at -100 mV was normalized as 1, and the normalized current in the absence or presence of inhibitor was picked up at same time point (**Fig. 3F**). A specific blocker of GIRK channels, TPN-Q(Jin *et al*., 1999), was used to confirm that the ACh-induced current was carried by GIRK channels. 3 µM TPN-Q completely suppressed the ACh-induced current and this effect was almost irreversible (**Fig. 3C**). The ACh-induced GIRK current was significantly inhibited by 10 µM and 100 μM BD1047 (**Fig. 3D-F**), showing that BD1047 also could inhibit the ACh-induced native GIRK current in isolated rat atrial myocytes.

**Figure 3.**
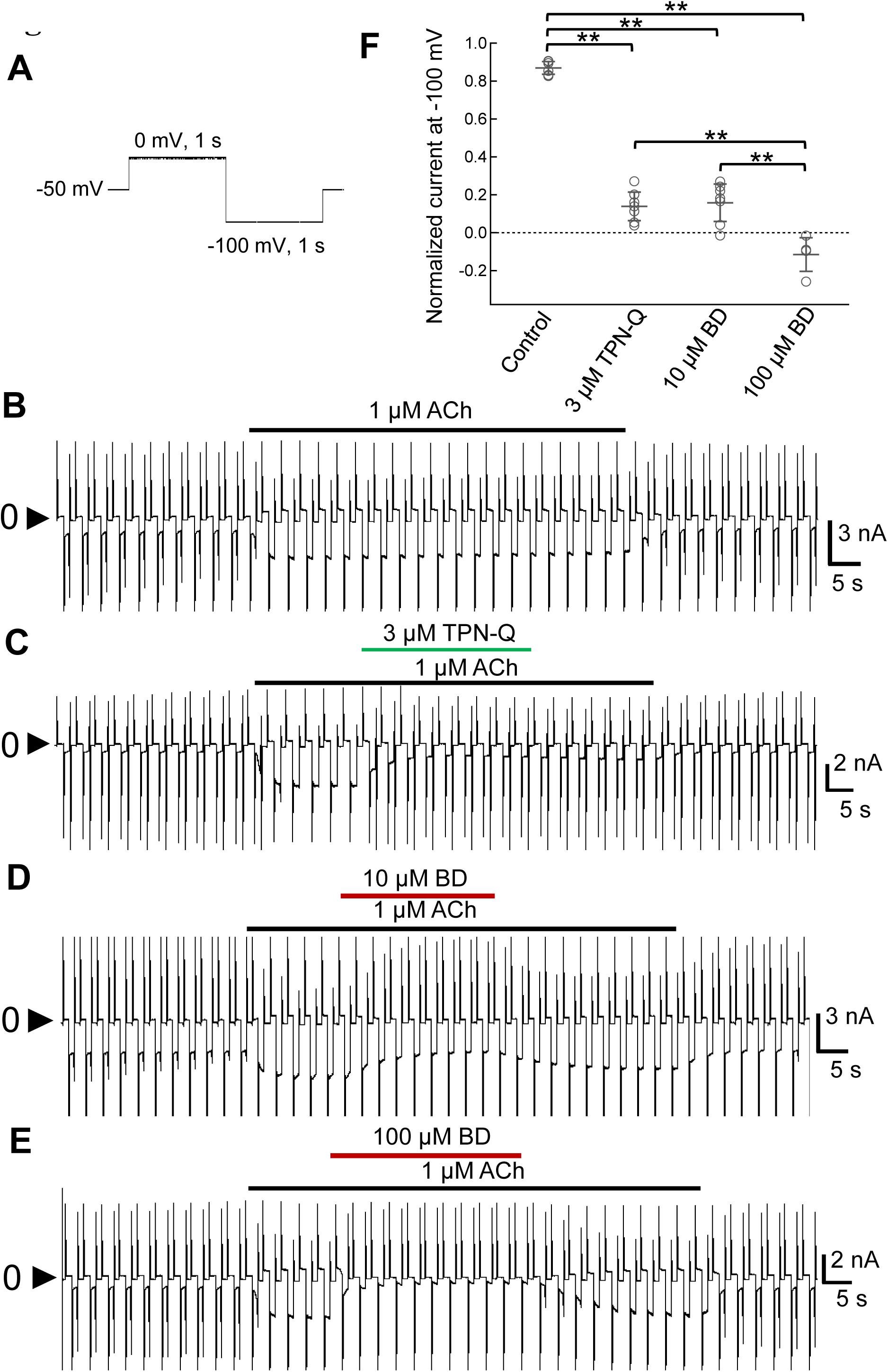
Effects of BD1047 on ACh-induced GIRK channel in rat atrial myocytes. **(A)** The voltage recording protocol used for patch-clamp recording from atrial myocytes. **(B-E)** The timelapse change of whole-cell currents recorded from the rat atrial myocytes in 20 mM K^+^ solution with **(B)** 1 μM ACh alone, **(C)** ACh with 3 μM TPN-Q (green bar), **(D)** ACh with 10 μM BD1047 (red bar) or **(E)** 100 μM of BD1047 (red bar). **(F)** Comparison of the normalized current after the application of inhibitor for 27 s. The control group indicates the desensitization of ACh-induced current which is calculated at the same time point as other groups. Data are mean ± SD (n = 5-7 for each); One way ANOVA followed by Tukey’s test, ** indicates *P*<0.01.

### Identification of the structural determinants of GIRK4 channel for the inhibition by BD1047

Taking advantage of the different sensitivities of GIRK2 and GIRK4 to BD1047, chimeras were generated to identify structural determinants of BD1047 inhibition. Five regions of GIRK2 (N-ter, transmembrane domain 1 (TM1), the pore forming loop between TM1 and TM2, TM2 and C-terminal (C-ter)) were substituted for the equivalent region of GIRK4 (**Fig. 4A**) to generate six chimeras, GIRK42222, GIRK24222, GIRK22422, GIRK22242, GIRK22224 and GIRK24242 **(Fig. 4A, 4B)**. The percentage inhibition by BD1047 for the chimeras containing the N-ter (GIRK42222), pore forming region (GIRK22422) or C-ter (GIRK22224) of GIRK4 were reached 61.8 ± 9.9 %, 37.1 ± 14.5 % and 49.2 ± 2.5 %, respectively compared to 19.7 ± 3.9 % for GIRK2 (**Fig. 4C**). This shows the importance of the N-ter, pore forming region or C-ter of GIRK4 for the inhibition by BD1047. The variation of the extent of inhibition of GIRK2 in Fig. 1F (37.7 %) and Fig. 4C (19.7 %) was judged to be due to the difference of batch of oocytes. Thus, we always included control group in each set of experiment.

**Figure 4.**
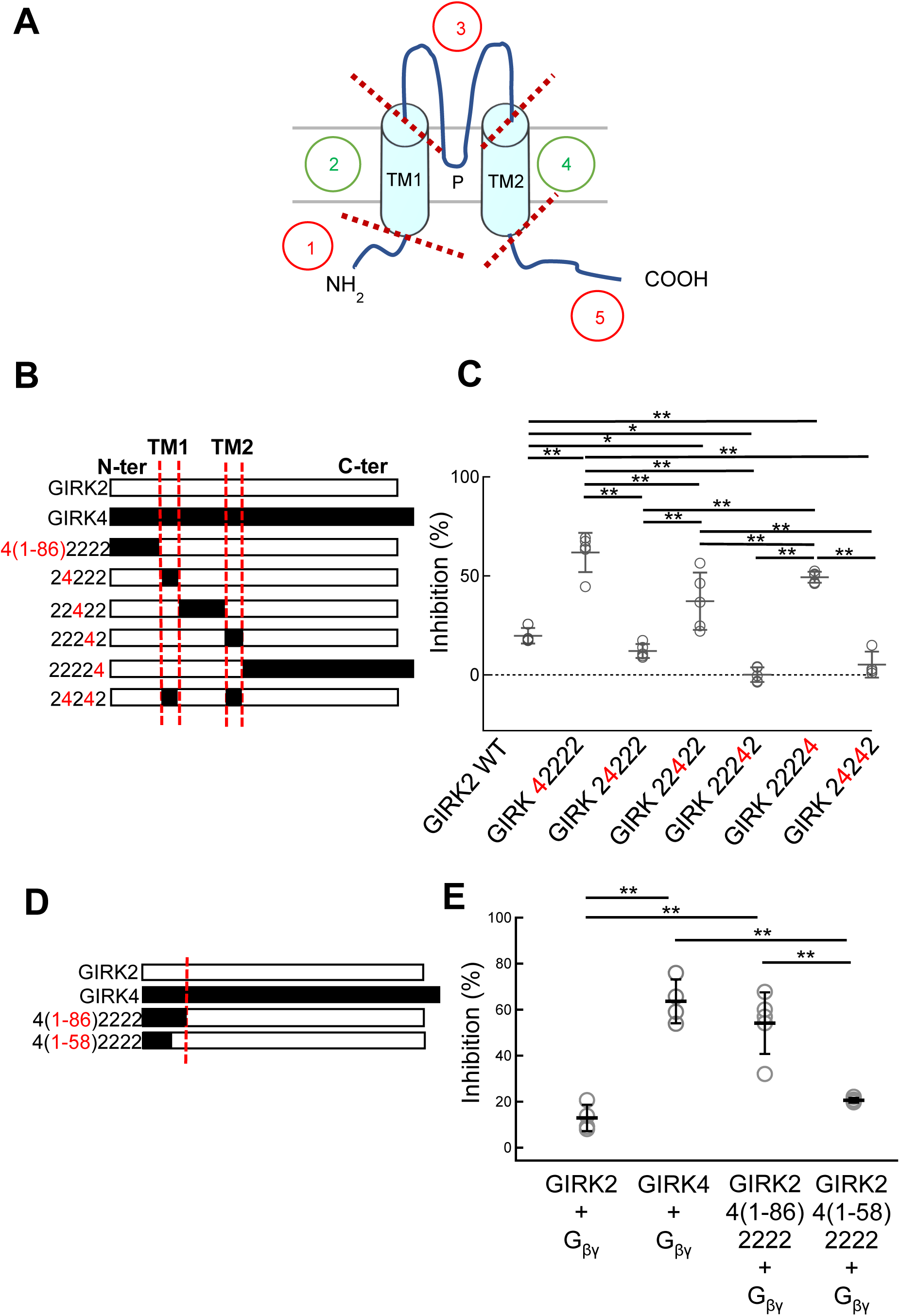
The effects of BD1047 on GIRK2/4 chimeras. **(A)** Chimeric constructs were generated in which 5 regions of GIRK2 (N-ter, TM1, the pore forming loop between TM1 and TM2, TM2 and C-ter) were substituted with the corresponding regions of GIRK4. **(B)** Schematic drawing of six chimeras, GIRK42222, GIRK24222, GIRK22422, GIRK22242, GIRK22224 and GIRK24242. Red letters “4” indicate those swapped to the corresponding part from GIRK4. **(C)** Comparison of the inhibition percentages before and after the application of 100 µM BD1047 to GIRK42222, GIRK24222, GIRK22422, GIRK22242, GIRK22224 and GIRK24242 chimeras. G_βγ_ subunits are co-expressed in all cases. **(D)** Schematic drawings of GIRK2/4 N-ter chimeras. GIRK2 4(1-86)2222 includes the whole 86 amino acids of N-ter from GIRK4, and GIRK2 4(1-58)2222 includes the distal 58 amino acids of N-ter from GIRK4. **(E)** Comparison of the inhibition percentages before and after the application of 100 µM BD1047. Data are mean ± SD (n = 4-5 for each); One way ANOVA followed by Tukey’s test, * indicates *P*<0.05, ** indicates *P*<0.01.

As the chimera GIRK42222 showed the strongest inhibition, a further chimera was generated in which only the most distal 58 amino acids (out of 86 within the N-terminus) were substituted (GIRK2 4(1-58)2222) (**Fig. 4D**). Inhibition by BD1047 of the chimera containing the distal region of N-ter (1-58) was only 20.6 ± 1.0 % (which is close to that of GIRK2 WT, 12.9 ± 5.7 %), while the inhibition ratio of the chimera containing the whole N-ter is 54.1 ± 13.4 % which is close to that of GIRK4 WT, 63.6 ± 9.5 % (**Fig. 4E**). These data show that the distal region of N-ter of GIRK4 is not involved in the inhibitory effect by BD1047 and highlights the importance of the proximal region of N-ter (59-86) of GIRK4.

To further narrow down the critical binding site of BD1047 on the proximal N-ter of GIRK4, single point mutants on GIRK2 and GIRK4 were made. These two channels differ by only three amino acids within the proximal N-terminus (**Fig. 5A**). For Arg73, Thr80 and Ile82 in GIRK2, the equivalent residues in GIRK4 are Gln, Ser and Leu respectively. These were individually mutated in GIRK2 and only GIRK2 I82L showed an increase in BD1047 sensitivity similar to GIRK4 (61.0 ± 1.9 % compared to 58.9 ± 3.0 %) (**Fig. 5B**). Consistent with this result, the reverse mutation in GIRK4 reduced inhibition by BD1047 (35.2 ± 6.6%) (**Fig. 5C**). Taken together, these results highlight the important of Leu77 in GIRK4 in determining the inhibitory effect of BD1047 on GIRK channels.

**Fig. 5.**
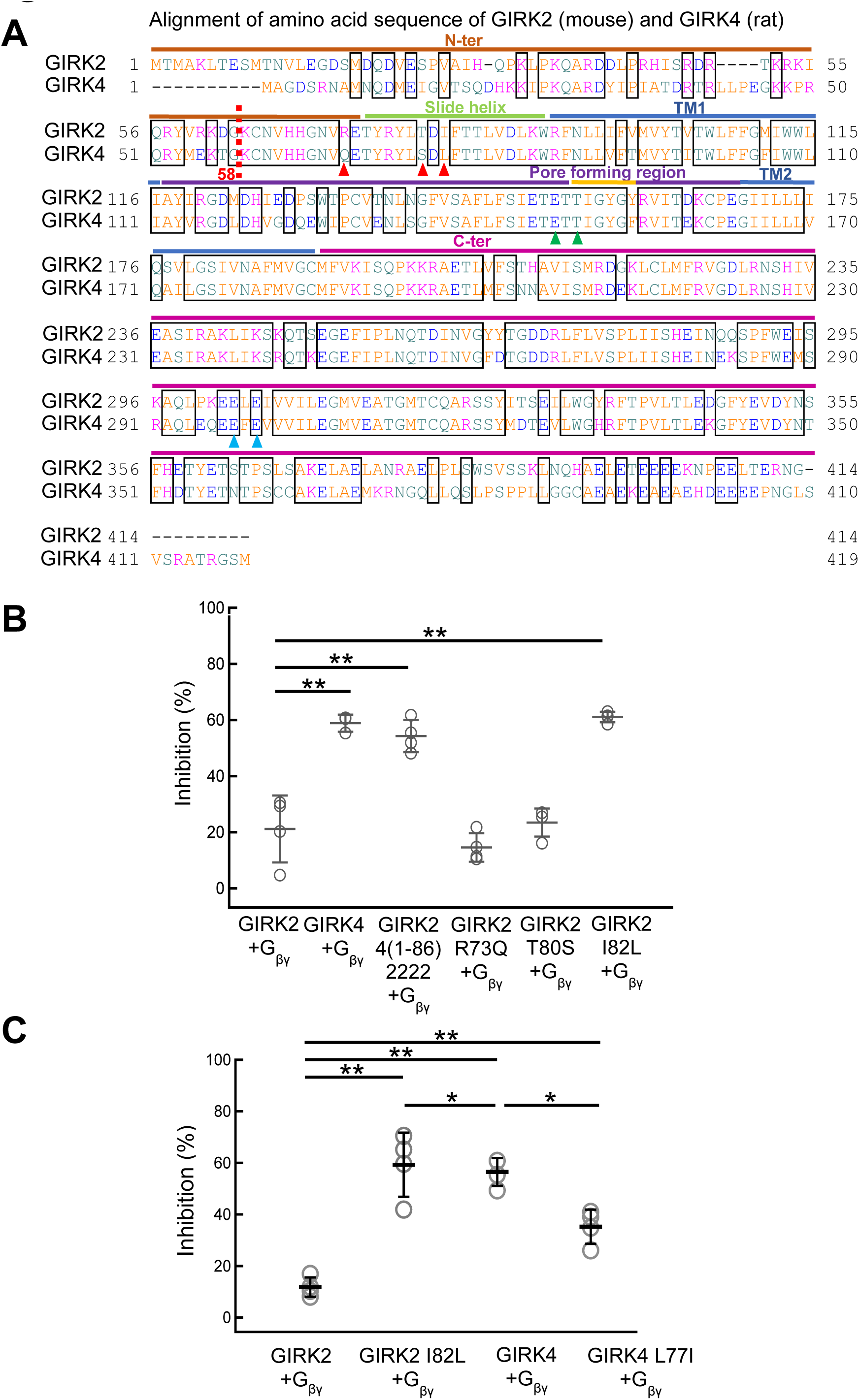
The effects of BD 1047 on GIRK2 and GIRK4 mutants. **(A)** The amino acid sequence alignment of mouse GIRK2 and rat GIRK4. A red dotted line indicates the start of the proximal N-ter of GIRK2 and GIRK4. Red arrows indicate non-conserved amino acids in the proximal N-ter region. Green arrows indicate the possible binding sites of BD1047 to GIRK4 on the pore forming region. Blue arrows indicate the possible binding sites between BD1047 and GIRK4 on the C-ter. **(B)** Comparison of the inhibition percentages before and after the application of 100 µM BD1047 on GIRK2, GIRK4, GIRK2 4(1-86)2222, GIRK2 R73Q, GIRK2 T80S and GIRK2 I82L. **(C)** Comparison of the inhibition percentages before and after the application of 100 µM BD1047 on GIRK2, GIRK2 I82L, GIRK4 and GIRK L77I. Data are mean ± SD (n = 4-5 for each); One way ANOVA followed by Tukey’s test, * indicates *P*<0.05; ** indicates *P*<0.01.

### Molecular docking of BD1047 and GIRK4 channel

Molecular docking is a useful approach to computationally predict the preferred binding site and docking orientation of a small molecule with a target protein. Thus, computational docking using the software named SwissDock online was conducted to identify the possible binding sites of BD1047 on GIRK4. As the structure of GIRK4 has not been solved yet, a homology structural model of GIRK4 was made based on the available structure of GIRK2 (PDB ID: 6XIT) using SWISS-MODEL online. Then, the obtained structure of GIRK4 and the chemical structure of BD1047 were uploaded to the SwissDock site.

The computational docking result of GIRK4 and BD1047 showed that BD1047 (cyan clusters) can bind to GIRK4 channel including N-ter, pore forming region as well as C-ter (**Fig. 6A**). Besides demonstrating the possible binding positions, the docking result also predicts all the possible binding directions and orientations of BD1047 at each predicted position so that BD1047 shown in cluster form. Therefore, when the intensity of the clusters at a position is higher, the possibility that BD1047 can bind to that position is judged to be higher. BD1047 clearly showed the highest intensity of clusters at the N-ter, indicating the highest chance that BD1047 interacts with GIRK4 at the N-ter (**Fig. 6A**). By highlighting the Leu77 residue on GIRK4 (**Fig. 6A**), it clearly demonstrates the importance of the Leu77 residue on GIRK4 for the binding with BD1047 which is consistent with the mutational analysis (**Fig. 5C**). On the contrary, the BD1047 cluster does not appear on the N-ter in the docking of BD1047 with GIRK2 channel (**Fig. 6B**). Interestingly, the docking of BD1047 on N-ter of GIRK4 is completely lost when Leu77 is mutated to Ile which is the corresponding amino acid on GIRK2 (**Fig. 6C**), whereas the cluster of BD1047 on the N-ter shows up in GIRK2 I82L mutation (**Fig. 6D**). Taken together, these results demonstrate the significant role of GIRK4 Leu77 for the binding of BD1047.

**Fig. 6.**
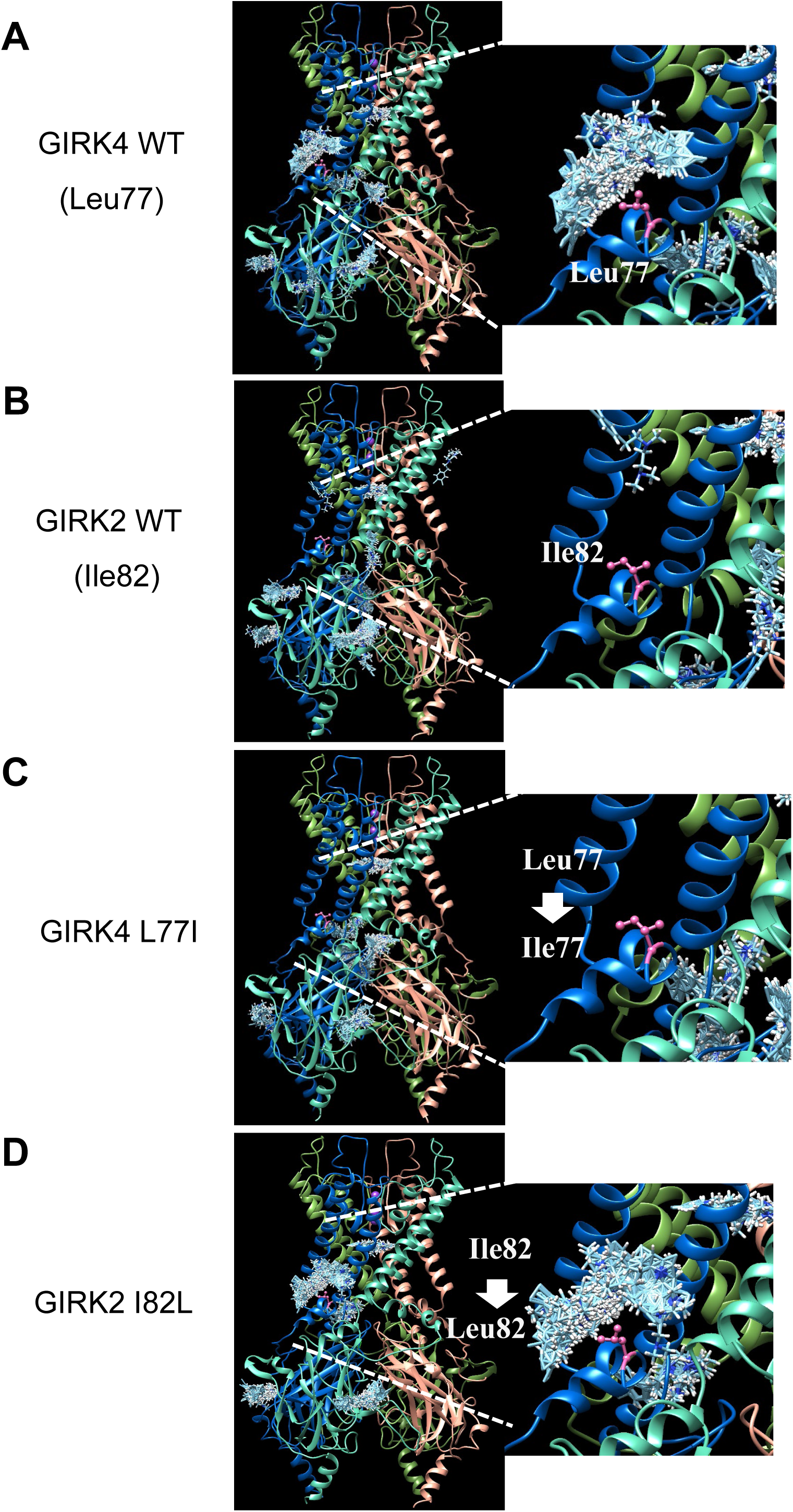
Computational molecular docking of BD1047 to GIRK4 and GIRK2. **(A-B)** The homology structure model of GIRK4 **(A)** and GIRK4 L77I **(B)** based on the structure of GIRK2 (6XIT). Cyan clusters of BD1047 indicate the predicted dockings of BD1047 to GIRK4 WT or GIRK4 L77I in various directions and orientations. Leu77 or Ile77 is highlighted in pink color. **(C-D)** The structure of GIRK2 (6XIT) **(C)** and homology structure model of GIRK2 I82L **(D)** based on the GIRK2. Cyan clusters of BD1047 indicate the predicted dockings of BD1047 to GIRK2 WT or GIRK2 I82L in various directions and orientations. Ile82 or Leu82 is highlighted in pink color. The right panels are an enlarged image of BD1047 docking on N-ter region in each left panel.

Meanwhile, there are some other amino acids adjacent to Leu77 in GIRK4, such as Tyr73, Leu74, Thr80, leu81 and Leu84, which might also play critical roles for the inhibition by BD1047, since it appears that they also interact with BD1047 (**Fig.7A**). These amino acids could not be identified by comparing the sequence alignment between GIRK2 and GIRK4, because they are conserved in both channels. We picked up some of them (Leu74, Leu81 and Leu84) and examined these point mutants. The inhibition effect of BD1047 was reduced for all of these mutants (**Fig. 7B**), thus providing support for the validity of the computational docking results.

**Fig. 7.**
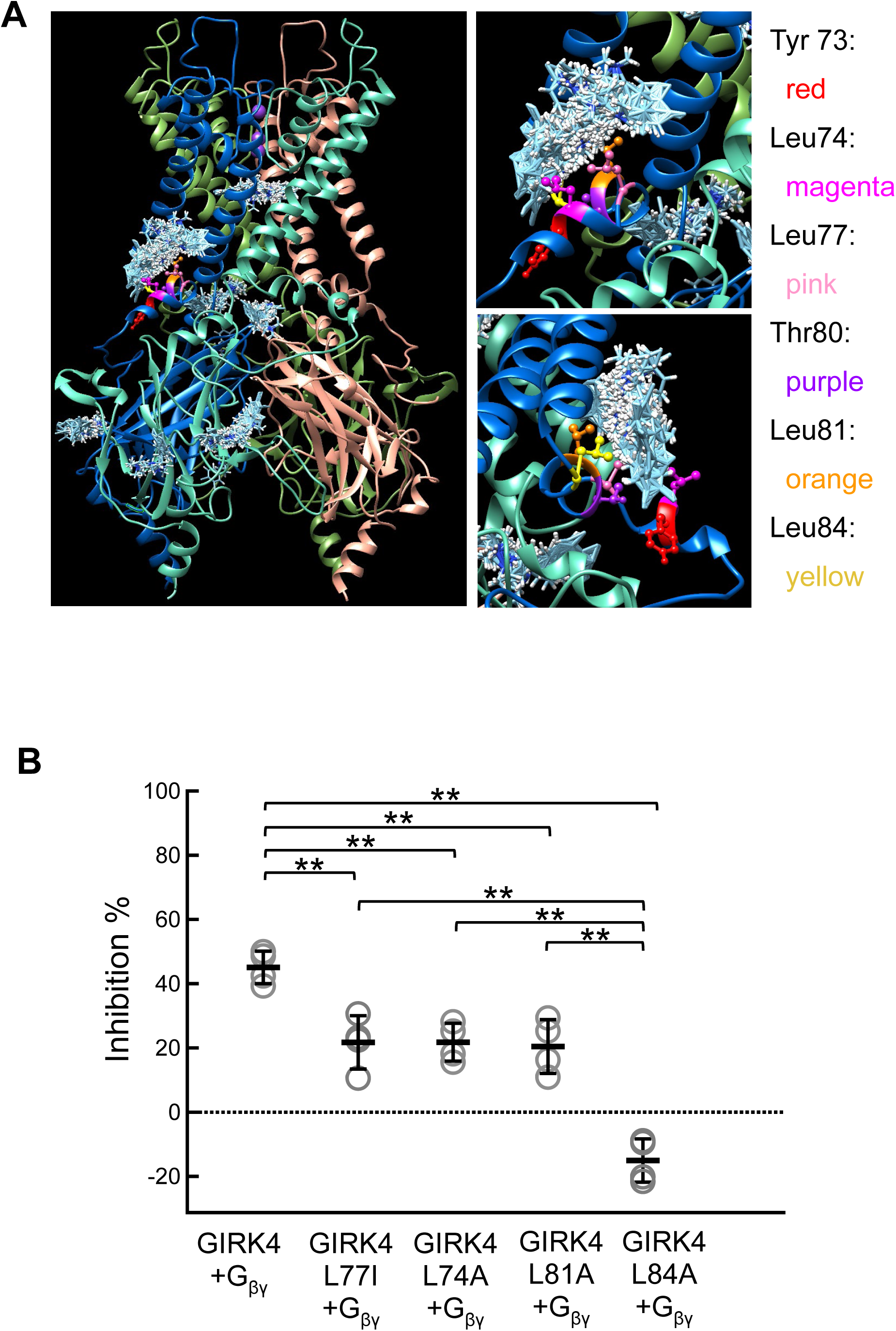
Computational molecular docking of GIRK4 and BD 1047 showing contribution of amino acid residues adjacent to Leu77. **(A)** The predicted protein structure is the homology model of GIRK4 based upon the structure of GIRK2 (6XIT). Cyan clusters of BD1047 indicate the predicted dockings in various directions and orientations of BD1047. Tyr73, Leu74, Leu77, Thr80, Leu81 and Lue84 are highlighted in red, magenta, pink, purple, orange and yellow colors, respectively. Right panels are enlarged images of Leu77 and adjacent amino acid residues for BD interaction from the left panel. **(B)** The effects of BD1047 on GIRK4 and its mutants of the predicted docking sites. Comparison of the inhibition percentages before and after the application of 100 µM BD1047 on oocytes expressing GIRK4, GIRK4 L77I, GIRK4 L74A, GIRK4 L81A and GIRK4 L84A. Data are mean ± SD (n = 4 for each); One way ANOVA followed by Tukey’s test, ** indicates *P*<0.01.

### The binding sites for BD1047 on GIRK4 involves the pore forming region and C-terminus

The analysis of the chimeras (**Fig. 4C**) and molecular docking analysis (**Fig. 6A, 7A**) indicated that not only the N-ter, but also the pore forming region and C-ter are involved in the inhibition of GIRK4 channel by BD1047. The residues Glu147 and Thr149 residues within the pore forming region **(Fig. 5A, 8A)** and Glu289 and Glu300 within the C-ter **(Fig. 5A, 8B)** were selected as candidates for further analysis since they showed a relatively high probability of interaction with BD1047 from the docking results. Alanine substitutions were made at each of the four positions. GIRK4 T149A and GIRK4 E300A channels appeared to be non-functional (**Fig. 8C**) whereas GIRK4 E298A was functional but there was no change in sensitivity to BD1047 compared to the wild type channel (**Fig. 8D**). In contrast, GIRK4 E147A showed a significant reduction in sensitivity to BD1047 (from 68.5 ± 7.2 % to 24.7 ± 7.7 %) (**Fig. 8D**). These data suggest that the Glu147 residue within the pore forming region of GIRK4 channel is also involved in the inhibition by BD1047.

**Fig. 8.**
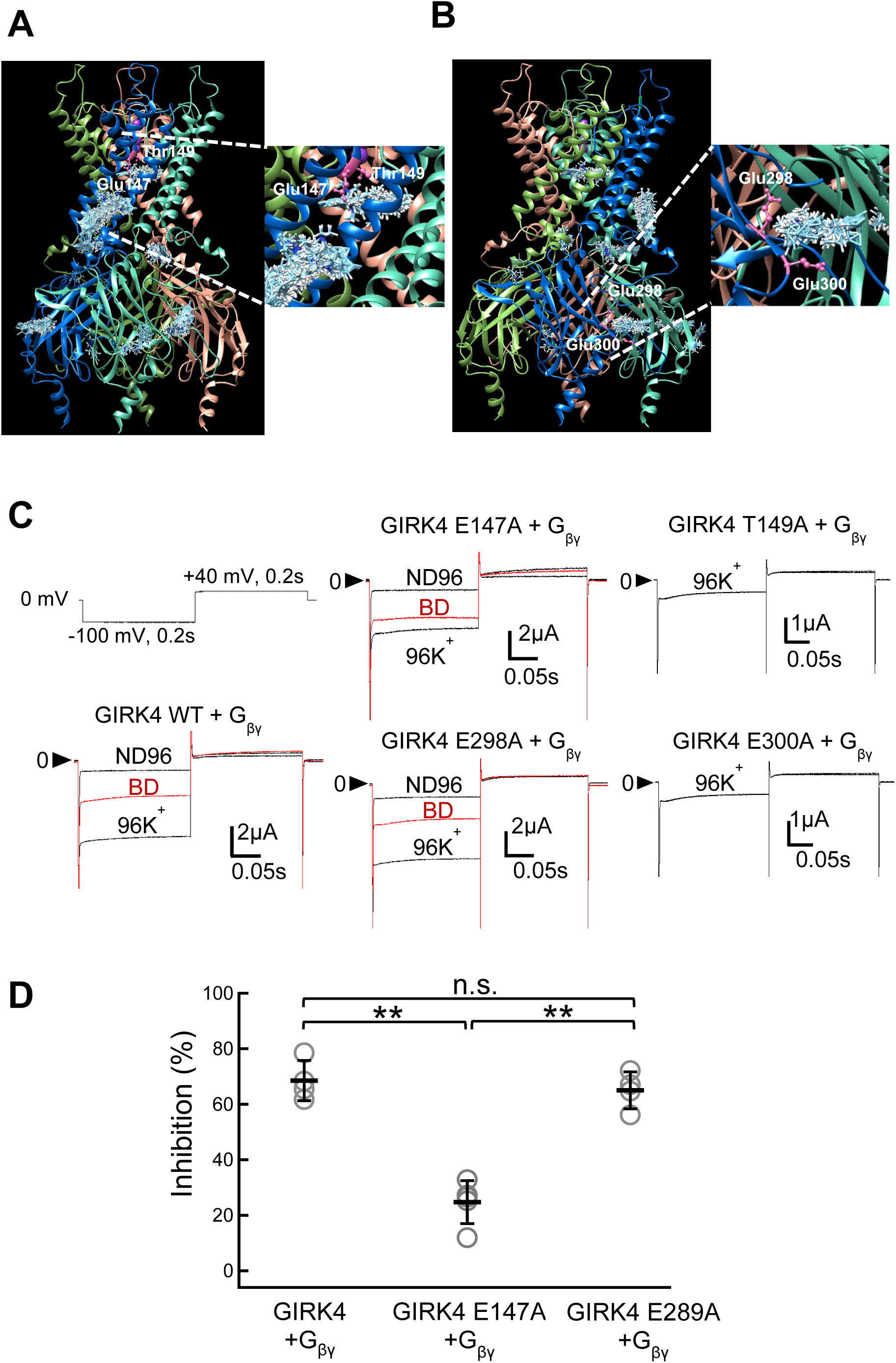
Identification of the binding sites of BD1047 to GIRK4 in the pore forming region and the C-ter. **(A-B)** Using the homology model of GIRK4 based on the structure of GIRK2 (6XIT), cyan clusters of BD1047 indicate the predicted dockings in various directions and orientations of BD1047. **(A)** The possible binding sites of BD1047 on the pore forming region of GIRK4. Glu174 and Thr149 are highlighted in pink. **(B)** The possible binding sites of BD1047 on the C-ter region of GIRK4. Glu298 and Glu300 are highlighted in pink. **(C)** Representative current recordings in ND96, 96K^+^, 96K^+^ with 100 μM BD1047 of GIRK4 WT, GIRK4 E147A, GIRK4 T149A, GIRK4 E289A and GIRK4 E300A in *Xenopus* oocytes evoked by the voltage protocol shown in the figure. G_βγ_ subunits are co-expressed in all five cases. **(D)** Comparison of the inhibition percentages before and after the application of 100 µM BD1047 on GIRK4 WT, GIRK4 E147A and GIRK4 E289A. Data are mean ± SD (n = 4-5 for each); One way ANOVA followed by Tukey’s test, ** indicates *P*<0.01.

### Competition between inhibitor and activator at Leu77 on GIRK4

IVM is an activator of GIRK channels, and a critical determinant of GIRK2 sensitivity to IVM is Ile82, which is equivalent to Leu77 in GIRK4 (Chen *et al*., 2017). Because this residue in involved in binding both IVM and BD1047 (**Fig. 9A**), the activator and inhibitor are likely to compete. To test this, we measured the concentration response relationship for BD1047 at GIRK4 in the absence and presence of 100 µM IVM (**Fig. 9B**). In the presence of IVM, there was a clear rightward shift in this relationship, with the IC_50_ of BD1047 in the presence of IVM of 62.9 ± 6.0 µM compared to 19.6 ± 0.5 µM in its absence (**Fig. 9B**). In addition, we compared the inhibition kinetics of GIRK4 and calculate the speed of the decrease of the GIRK4 current upon the application of BD1047 in the absence and presence of IVM (**Fig. 9C, 9D**). The prensence of IVM right shifted the BD1047’ inhibition kinetics (**Fig. 9E**), the tau of inhibition in the absence and presence of IVM are 11.8 ± 1.1 s and 22.4 ± 1.9 s respectively (**Fig. 9F**). The data demonstrates a slowing of BD1047’ inhibition of GIRK4 channels by IVM. In conlusion, these results suggest that IVM could affect the binding of BD1047 and compete with BD1047 at Leu77 on GIRK4.

**Fig. 9.**
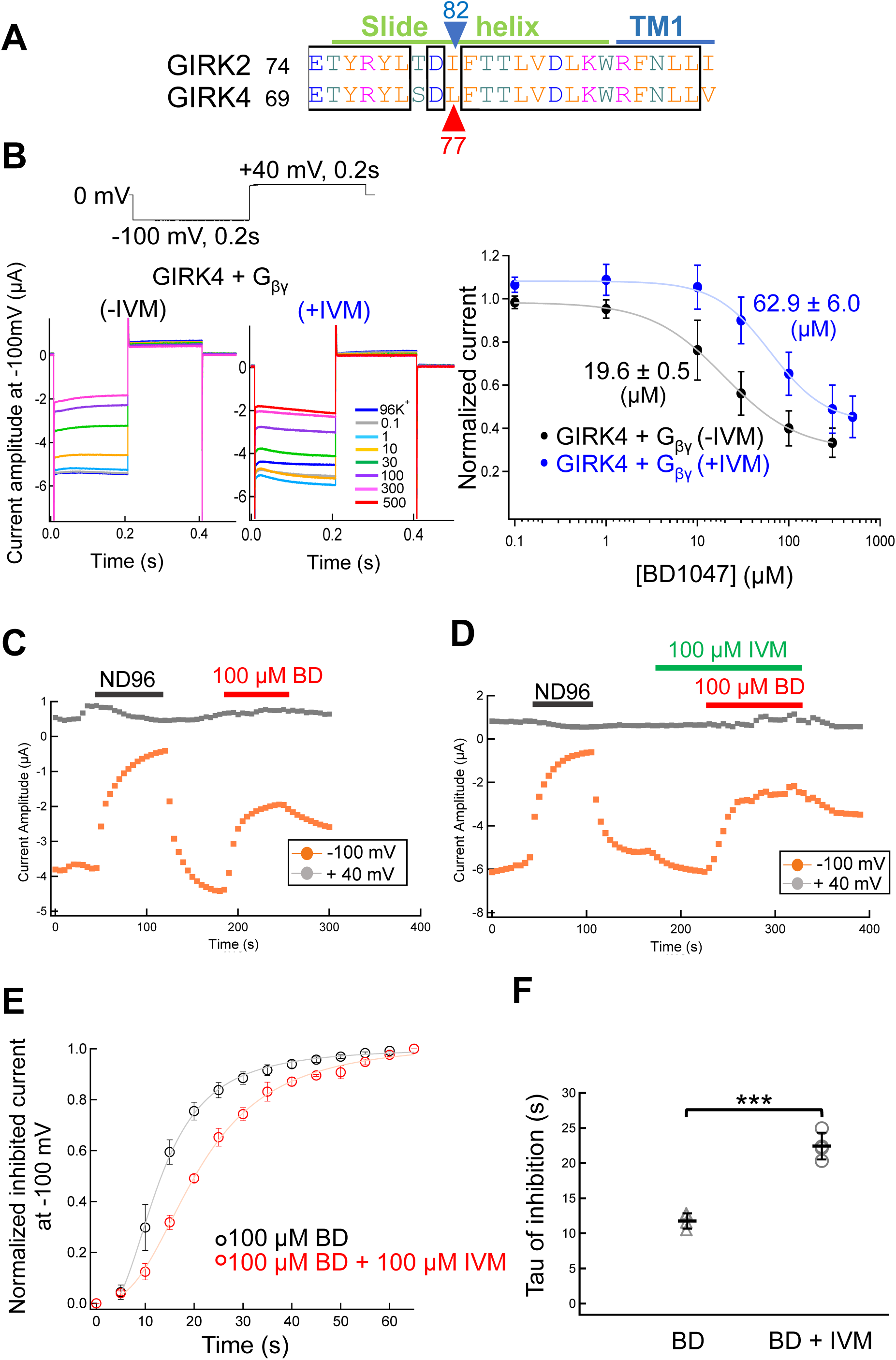
Competition between BD1047 and IVM on GIRK4. **(A)** The amino acid sequence alignment of mouse GIRK2 and rat GIRK4. The blue arrow and red arrow indicate the Ile82 of GIRK2 and the corresponding amino acid Leu77 on GIRK4, respectively. **(B)** The effects of 100 μM IVM on the concentration-response relationship of BD1047 on GIRK4 channel. Left and middle: the representative current recordings in *Xenopus* oocytes. Various concentration of BD1047 from 0.1 μM to 500 μM were applied to GIRK4 channel in the absence of IVM (Left) and presence of IVM (Middle). Right: Dose-inhibition relationships of BD1047 on GIRK4 channel in the absence of IVM (black) and presence of IVM (blue). Data are mean ± SD (n = 4-5) for each plot. IC_50_ is 19.6 ± 0.5 (μM) (-IVM) and 62.9 ± 0.5 (μM) (+IVM), respectively. **(C)** The timelapse changes of the current amplitudes at -100 mV (orange plots) in 96K^+,^ ND96 and 96K^+^ with 100 µM BD1047 solution in oocytes only expressing GIRK4+G_βγ_ channels. **(D)** The timelapse changes of the current amplitudes at -100 mV (orange plots) in 96K^+^, ND96 (black bar), 96K^+^ with 100 µM IVM (green bar), 96K^+^ with 100 µM BD1047 (red bar) and IVM solution (green bar). **(E)** Effects of IVM on the inhibition kinetics of BD1047 on GIRK4 current. The inhibition kinetics calculated from the current amplitude at −100 mV after the application of BD1047 in the absence or presence of IVM. The maximal reduction of current after the application of BD1047 was normalized as 1. **(F)** Tau of BD’ inhibition calculated from (E). Data are mean ± SD (n = 4 for each); unpaired *t*-test, *** indicates P<0.001.

## Discussion

In this study we investigated the mechanism of action of BD1047 at GIRK channels and showed (1) BD1047 directly inhibits GIRK channel currents independent of the sigma-1 receptor and produces a greater inhibition of GIRK4 compared to GIRK2; (2) It also inhibits ACh-induced native GIRK current in isolated rat atrial myocytes; (3) Switching the proximal cytoplasmic N-terminal regions of GIRK2 and GIRK4 is sufficient to switch sensitivity to BD1047 and within this region of GIRK4, Leu77 is a critical determinant of this inhibition; (4) Molecular docking analysis similarly suggests that Leu77 is contributing to the BD1047 binding site on GIRK4; (5) An activator of GIRK channels, IVM, competes with BD1047 at Leu77 on GIRK4.

### The antagonism of Sigma-1 receptor by BD1047

BD1047 was identified as an antagonist of S1R by binding assays and behavioral studies in rat. In radioligand binding studies using rat liver and guinea pig brain, BD1047 bound S1R with high affinity. In rats, the microinjection of BD1047 into the red nucleus decreased dystonia induced by haloperidol, a known S1R agonist (Matsumoto *et al*., 1995). In other studies, a possible usage of BD1047 was suggested in alcohol use disorder because BD1047 could attenuate ethanol-induced intracellular Ca^2+^ in rat hippocampal through the modulation of the function of S1R and ER-bound IP_3_ (Reynolds *et al*., 2016). In a recent study, BD1047 was shown to block intracellular calcium responses in cultured astrocytes and to alleviate S1R-induced pain hypersensitivity in naïve mice (Choi *et al*., 2022). There has been an assumption that BD1047 is a selective antagonist at S1R, however, our studies show that BD1047 can also affect other molecules such as GIRK channels in addition to S1R.

### The mechanism of the binding of BD1047 to GIRK4 channel

The mutagenesis studies highlight L77 as playing an important role in BD1047 mediated inhibition of GIRK4, however BD1047 is still more potent at GIRK4 L77I compared to GIRK2 WT (**Fig. 5C**) suggesting that other region(s) of GIRK4 contribute to the binding of BD1047. In the analysis of concentration-response relationship of BD1047, the voltage-dependent slow activation upon hyperpolarization of GIRK2 channel change was observed only in the presence of high concentration of BD1047 (**Fig. 2A**). The change of the kinetics of the voltage-dependent activation may represent multiple binding sites of BD1047 to GIRK2 channel. For example, BD1047 only binds to high affinity binding site on GIRK2 when the concentration is low, while it also binds to low affinity binding site to induce slow activation when the concentration is high (**Fig. 2A**). In addition, GIRK2/4 chimera studies also showed a possible involvement of the pore forming region and the C-ter of GIRK4 (**Fig. 4C**), which are consistent with the results of the molecular docking study (**Fig. 6A**). Meanwhile docking data indicated some amino acids residues which are conserved between GIRK2 and GIRK4 on the proximal N-ter also have high chance to interact with BD1047 (**Fig. 7A**). By performing mutagenesis studies on the N-ter, pore forming region and C-ter of GIRK4, we identified further more amino acid residues, Leu74, Leu81, Leu84 in the N-ter and Glu147 in the pore region, which are also involved in the inhibition by BD1047 **(Fig. 7B, 8D)**. These results probably help us explain why there was only partial decrease in the BD1047’s inhibition when just one single amino acid, Leu77, was mutated on GIRK4.

Interestingly, the docking of GIRK4 L77I and BD1047 showed a loss of the N-ter cluster of BD1047, whereas GIRK2 I82L showed an acquisition of it. Leucine and isoleucine are both hydrophobic amino acids in the same category. The structural difference between them is that the leucine contains an isobutyl side chain with two of methyl in the end to form a mermaid-tail-like structure (**Fig. 10A**), while isoleucine contains a secbutyl side chain with one methyl in the end (**Fig. 10B**). By superposing the homology model of GIRK4 L77I with the structure of GIRK2 WT, it was confirmed that the side chain position and direction of Ile77 substituted into GIRK4 is the same as Ile82 in GIRK2 (**Fig. 10C**) and vice versa for GIRK2 I82L and GIRK4 (GIRK4 Leu77) (**Fig. 10D**). This comparison may explain the loss and gain of the docking of BD1047 after mutation on the N-ter of GIRK2 or GIRK4. Meanwhile, it seems that the mermaid-tail-like structure formed by two methyl in isobutyl side chain of leucine is very critical to form the binding pocket for BD1047.

**Fig. 10.**
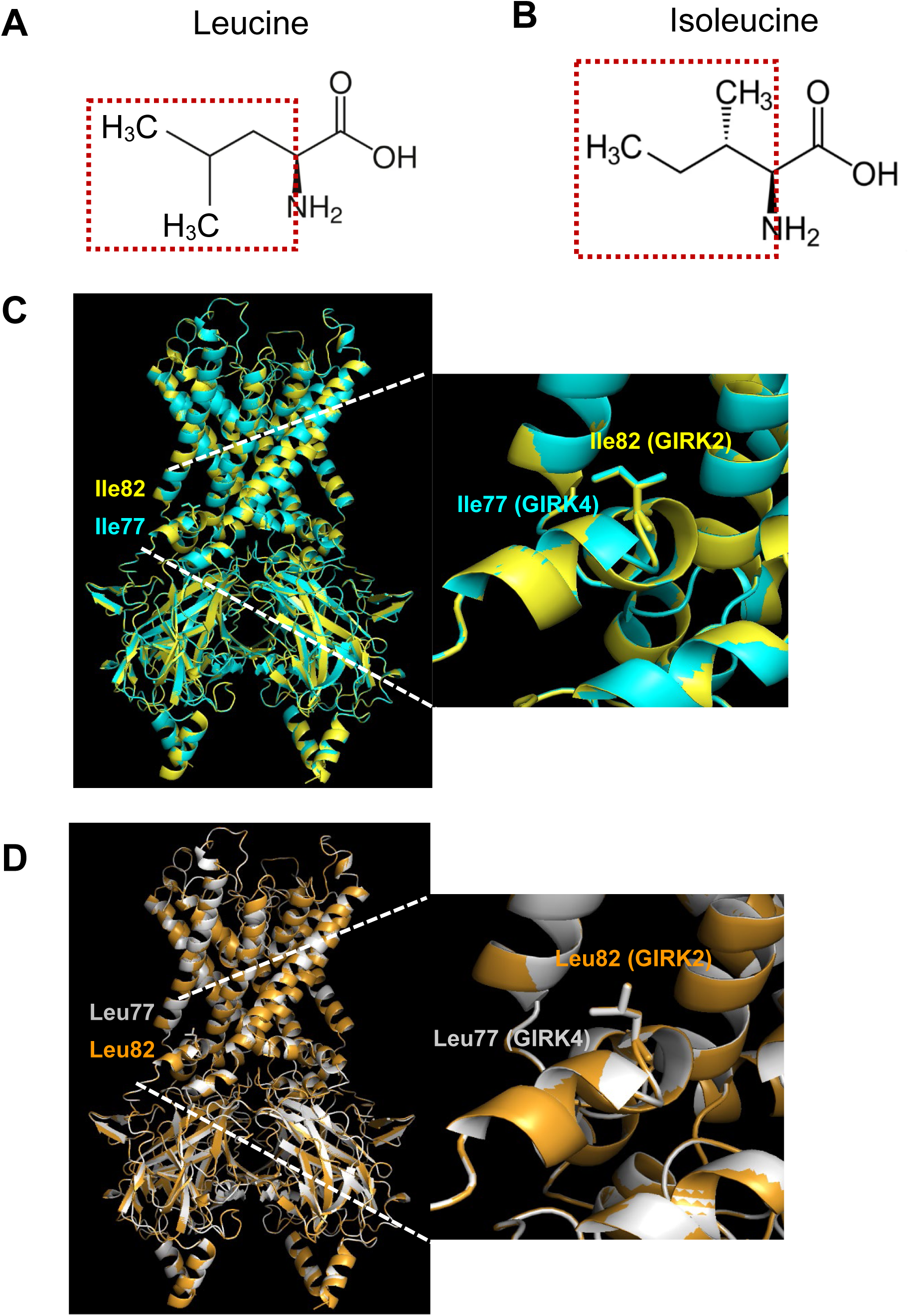
Comparison of the side chain between GIRK4 and GIRK2. **(A)** The chemical structure of leucine. Isobutyl side chain is highlighted with red frame. **(B)** The chemical structure of isoleucine. Secbutyl side chain is highlighted with red frame. **(C)** Left: The overlay of the structure of GIRK2 WT (6XIT) (yellow color) and homology model of GIRK4 L77I based on GIRK2 (6XIT) (cyan color). Right: an enlarged image from left panel. **(D)** Left: The overlay of the structure of homology model of GIRK4 WT based on GIRK2 (6XIT) (gray color) and homology model of GIRK2 I82L based on GIRK2 (6XIT) (orange color). Right: an enlarged image from left panel.

The results above show that computational docking data indeed help the confirmation of the incomplete mutagenesis study and provides quite comprehensive and accurate information. However, as its trustability is not necessarily guaranteed, the combined usage with electrophysiological analysis of chimeras and mutants performed in this study is indispensable to draw a solid conclusion. For example, docking data also demonstrated binding of BD1047 within the pore region and C-ter of GIRK4 which is consistent with chimera results, suggesting there might be some other essential amino acid residues for the inhibition. We tried some more mutagenesis study according to the docking results as well **(Fig. 8A, 8B)**. Glu147 and Thr149 on pore forming region as well as Glu298 and Glu300 on the C-ter has been mutated to Ala respectively, but the attenuation of the inhibition of BD1047 was only observed in GIRK4 E147A, not others (**Fig. 8C, 8D**).

### The Leu77 on GIRK4 may be a common drug binding site

Our group reported previously that the antiparasitic drug, IVM, can activate GIRK channels and the critical amino acid residue for the activation of the GIRK2 by IVM is Ile82. Ile82 is located in the slide helix between the TM1 and the N-terminal cytoplasmic tail domain (CTD) (Chen *et al*., 2017). In our experiments, Leu77 of GIRK4, correspondingly to Ile82 of GIRK2, was identified as the critical amino acid residue for the inhibition effect by BD1047. Furthermore, another group recently showed a role for GIRK4 Leu77 (corresponding to GIRK2 Ile82) in the activation of GIRK channels by a novel activator, 3hi2one-G4. The 3hi2one-G4 selectively activates GIRK4 channel via Leu77 and the effects of 3hi2one-G4 is significantly decreased when Leu77 is mutated to Ile77. (Cui *et al*., 2022). Taken together, both activator and inhibitor with different chemical structures bind to this slide helix region between TM1 and CTD of GIRK channel, suggesting that this position might be a shared hot spot for drug binding. Thus, it could be used as therapeutic target to develop new medicines for GIRK channel-associated diseases.

It has been reported that GIRK4 may play an important role in the heart arrhythmia because the GIRK1/4 channel is constitutively activated in the atrial myocytes of chronic AF patients (Dobrev *et al*., 2005) and in GIRK4 knockout mice, ACh did not induce AF (Kovoor *et al*., 2001). Thereby, it is beneficial to identify inhibitor(s) targeting GIRK4 channel as potential therapies for GIRK4-associated arrhythmia. Various inhibitors of GIRK channels such as tertiapin-Q, clozapine, fluoxetine or terfenadine have been identified, but they did not show selectivity towards homomeric GIRK4 channels (Kobayashi *et al*., 2000; Kanjhan *et al*., 2005; Kobayashi *et al*., 2011; Chen *et al*., 2019; Jeremic *et al*., 2021). Our results demonstrating that BD1047 is more potent at GIRK4 may help to optimize development of novel therapies for GIRK4-associated arrhythmia with reduced side effects caused by actions on other GIRK channels.

## Author contributions

Oocyte preparations and all of the electrophysiological experiments were performed in the laboratory of Y.K. in the division of Biophysics and Neurobiology, National Institute for Physiological Sciences (NIPS). C.L. and Y.K. designed the study. C.L. performed all mutagenesis and electrophysiological recordings using *Xenopus* oocytes and data analyses; I-S.C. performed isolation of atrial myocytes from rat and construction of GIRK2-GIRK4 chimers. M.T. performed electrophysiological recordings from atrial myocytes. C.L. and Y.K. wrote the paper. All authors have revised and approved the final version of the manuscript.

## Acknowledgments

The authors thank Ms. Naito, C. and Yamamoto, T. for technical assistance, and all members in Kubo laboratory for discussion. The authors also thank Dr. Ruth Murrell-Lagnado (Sussex University, School of Life Sciences, UK) for editing the manuscript.

